# A microfluidic-based model of nociceptor sensitization reveals a direct activation of sensory axons by prostaglandin E2

**DOI:** 10.1101/2022.03.18.484883

**Authors:** Georgios Kimourtzis, Natasha Rangwani, Bethan J. Jenkins, Peter A. McNaughton, Ramin Raouf

## Abstract

Prostaglandin E2 (PGE2) is one of the major contributors to inflammatory pain hyperalgesia, however, the extent to which it modulates the activity of the nociceptive axons is incompletely understood. We used a microfluidic cell culture platform to investigate the changes in responsiveness of sensory axons following treatment with PGE2. We show that the application of PGE2 to fluidically isolated axons leads to sensitization of their responses to depolarising stimuli, and the inclusion of zatebradine, a blocker of HCN channels, blocks this enhancement. However, unexpectedly, we also found that the application of PGE2 to the axons elicited a direct and persistent spiking activity in the sensory neurons. We demonstrate that this persistent activity is due to a direct depolarization of axons by PGE2, which is inhibited by Nav1.8 sodium channel blockers but is mainly refractory to Nav1.7 channel blockade. Both the persistent activity and the membrane depolarization in the axons are abolished by the EP4 receptor inhibitor and a blocker of cAMP synthesis. Our data indicate that PGE2/EP4/cAMP pathway culminates in a sustained depolarization in the sensory axons, leading to the generation of action potentials propagating to the soma. PGE2 therefore, not only mediates nociceptor sensitization but can directly elicit discharges in nociceptive axons, hence redefining its role as a pain mediator in inflammatory conditions.

**One Sentence Summary:** Prostaglandin E2 can depolarise nociceptive axons in the absence of any noxious stimuli leading to a sustained activation of pain sensing neurons.

## INTRODUCTION

Despite the recent advances, but primarily due to the complexity of the pain pathways, clinical management of pain remains an area of significant unmet need (*1*). PGE2, which is involved in the regulation of a number of processes in the nervous system, plays a central role in both inflammatory and neuropathic pain (see (*2*) and (*3*) for review). PGE2 belongs to a class of abundantly expressed eicosanoids, produced by enzymatic catalysis of arachidonic acid. Cyclooxygenases COX-1 and COX-2 produce the prostaglandin precursors PGG2 and PGH2 which are in turn converted by prostaglandin synthases into the active prostaglandins E2 (PGE2), D2 and F2A, prostacyclin (PGI2) and thromboxane (see (*4*) for review). The COX-2 inhibitors, which belong to a class of non-steroidal anti-inflammatory drugs (NSAIDS), are extensively used as analgesic compounds. However, the adverse effects, such as peptic ulcers and increased cardiovascular risks, associated with these inhibitors have made the development of new anti-inflammatory analgesics an area of urgent need (*5*).

PGE2 has been shown to bind to EP receptor subtypes (EP1-4), which are G-protein coupled receptors expressed in many tissues including the sensory neurons (*6*). The high affinity receptor for PGE2, EP4, forms a Gs complex, and activates adenylate cyclase which in turn increases cyclin AMP (cAMP) production **(*7, 8*)**. The PGE2/EP4 pathway has been shown to play an important role in osteoarthritis and rheumatoid arthritis **(e.g. Ref (*9*))**. In other animal models with CFA and carrageenan treatment, pharmacological antagonism of EP4, the high-affinity binding receptor for PGE2, was efficacious in attenuating the associated inflammatory hyperalgesia (*10, 11*).

The pronociceptive effects of PGE_2_ and its role in peripheral sensitization have been established in animal models and are supported by studies that show the sensitization of the nociceptive nerve endings in the skin and membrane targets by PGE2/EP4 pathway (see (*2*) and (*3*) for review). Although sensitization of sensory stimuli by PGE2 is well established, whether PGE2 can directly excite the nociceptive fibers remains incompletely understood. For instance, initial studies had suggested the excitation of nociceptors by prostaglandins (*12*), whereas others have reported that PGE2 fails to evoke spike discharge in visceral afferents and skin-saphenous nerve preparations (e.g.(*13, 14*)). Studies using conventional cell cultures have also been inconclusive. Some reports have shown that PGE2 evoked Ca2+ influx in dissociated rat DRG neurons (e.g. (*15*)), whereas others have reported that exogenous application of the same concentration of PGE2 did not elicit an increase in intracellular Ca2+ levels in DRG cell bodies (e.g. (*16*))

We have previously reported the development of a microfluidic culture platform (*17*) to investigate the function of sensory axons in isolation from the soma, as a cell culture model of nociceptive endings. We have shown that this model can recapitulate salient features of the physiology and anatomy the nociceptors, and is ideal for studying the functional changes in nociceptive axons in response to mediators such as NGF (*17, 18*). In the present study we investigate the modulation of axonal excitation in response to PGE2 using a microfluidic culture platform. We demonstrate that local treatment of axons with PGE2 in microfluidic cultures leads to sensitization of the responses to depolarising stimuli, but also induces a persistent activity in many nociceptive axons by causing a direct depolarization of the axons in the absence any other stimuli. We provide evidence for the ionic mechanism of PGE2-evoked excitation of nociceptive axons and show that Nav1.8 channels are indispensable for the depolarization and the concomitant persistent activity in sensory axons.

## RESULTS

### PGE2 sensitizes axonal responses to KCl in the microfluidic cultures

In order to investigate PGE2 modulation of axonal function, we employed a microfluidic culture platform. We had previously shown the versatility of this culture platform for examining axonal responses to various stimuli (*17*). As demonstrated by Tsantoulas et al. (2013), chemical or electrical stimulation of fluidically isolated DRG axons results in a local depolarization and generation of action potentials that propagate to the cell soma. This triggers the opening of voltage-dependent calcium channels, resulting in an increase in intracellular Ca2+ at the soma (*17, 18*).

We examined the modulation of axonal responses to 20mM KCl by PGE2 using microfluidic cultures (**Figure1A)** where the somal compartment is fluidically isolated from the axonal compartment (*17*). The magnitude of the responses in the fluidically isolated DRG axons were evaluated by measuring the changes in intracellular Ca2+ at the soma as a readout for axonal function(*17*). Only DiI positive cell bodies, indicative of axonal crossings (see Methods), were selected for further analysis (**Figure S1**). Administration of KCl to axons led to a robust increase in Ca2+ concentration measured at the soma (**Figure 1B**, and **Figure S2**).

**Figure 1.**
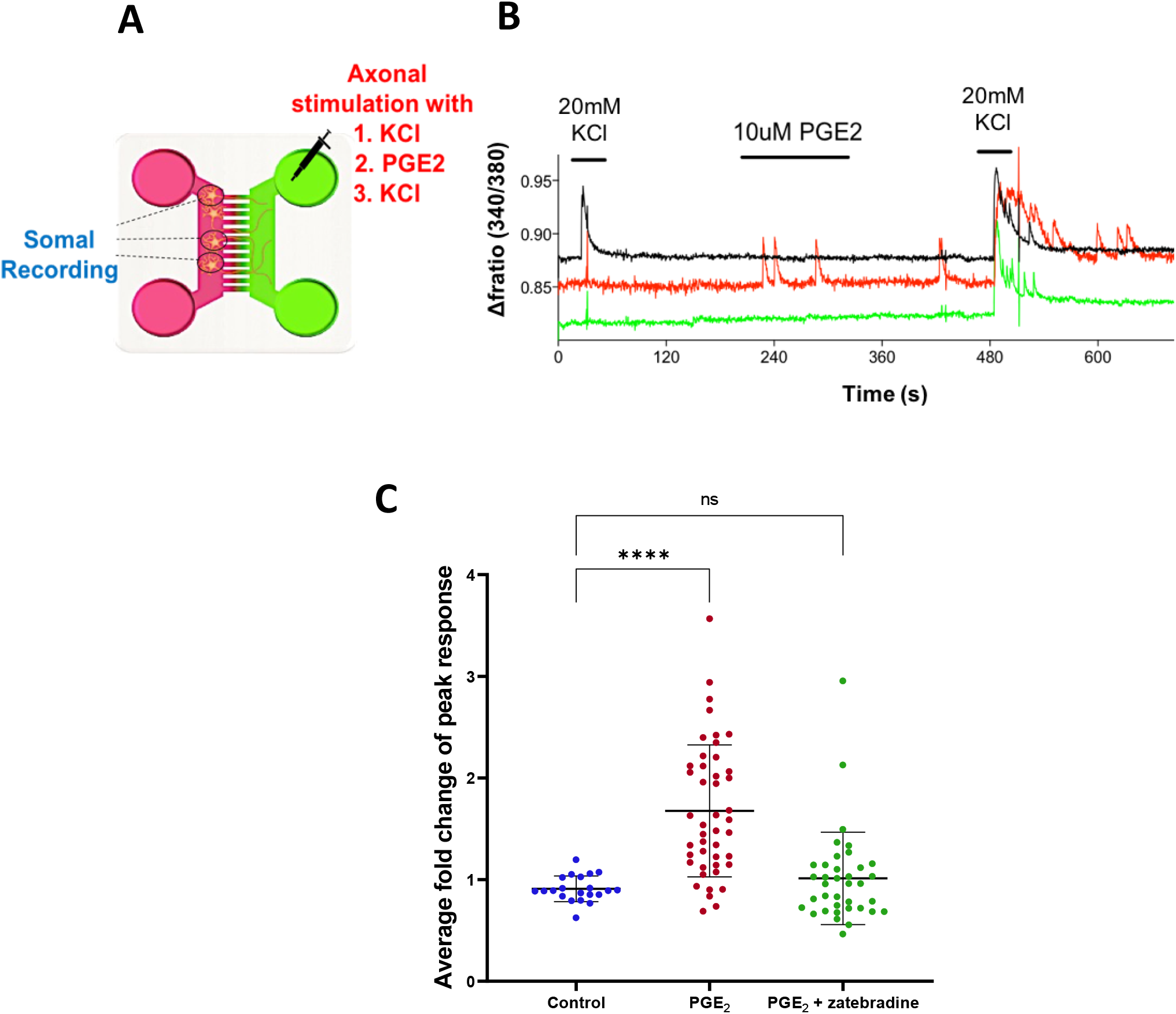
PGE2-mediated sensitization of response to KCl stimulation of axonal compartment in microfluidic cultures. A) Schematic of the experimental setup using microfluidic cultures. Somal side (pink) is separated from the axonal side (green) by 150µm microgrooves. B) Representative Fura-2 traces recorded from the somal compartment while stimulating the axonal compartment with 20 mM KCl and a two-minute treatment with PGE2. C) Summary of the changes in peak responses to KCl stimulation after treatment with imaging buffer (control) or PGE2 and the effect of zatebradine. The potentiation of KCl-evoked axonal responses was abolished by inclusion of 10 mM zatebradine (****p<0.0001,Oneway ANOVA followed by Tukey’s t-test, ns p=0.2061, n=21 – 45 from 5 cultures).

PGE2 treatment of the axons led to the potentiation of the responses to KCl administered to the axons (**Figure 1B and C)**. This potentiation was observed using a a cocktail of inflammatory mediators (**Figure S3**).

The magnitude of Ca^2+^ responses displayed a 2.7-fold increase, from 0.88 ± 0.03 to 2.38 ± 0.12, to KCl stimulation following application of PGE2 (**Figure 1C)**. HCN channels have been shown to play a crucial role in PGE2 mediated pain sensitization in mice(*19, 20*). We investigated the effect of zatebradine, a blocker that is selective for HCN channels but does not discriminate amongst the isoforms HCN1-4, on the PGE2 mediated enhancement of axonal responses. We found that pretreatment of axons with zatebradine completely abolished the PGE2 mediated enhancement of KCl responses **(Figure 1D)**. Our results demonstrate that PGE2 sensitizes axonal responses to depolarising stimuli, and this enhancement is dependent on HCN channels.

### PGE2 evokes persistent firing activity in sensory axons

We had observed that the application of PGE2 after KCl stimulation of fluidically isolated sensory axons induced short, repetitive bursts of intracellular Ca^2+^ detected at the soma **(Figure 1B)**. We sought to determine whether PGE2 on its own could evoke firing activity in these sensory axons. We employed a three-compartment microfluidic device configuration where sensory axons from the neurons plated in the somal compartment, traversed the middle and distal compartments (**Figure 2A)**. A two minute application of PGE2 to the distal compartment produced a persistent axonal activity in the form of calcium transients detected at the soma compartment (**Figure 2B)**. We found that the ongoing activity continued following washout of PGE2. This persistent activity in the form of calcium transients was quantified as the total number of transients observed over 3 minutes following PGE2 application (**Figure 2E)**. PGE2 application to the distal axonal compartment produced transients at a rate of 6.12 ± 0.63 / 3 minutes after application, compared to no transients observed in the absence of PGE2.

**Figure 2.**
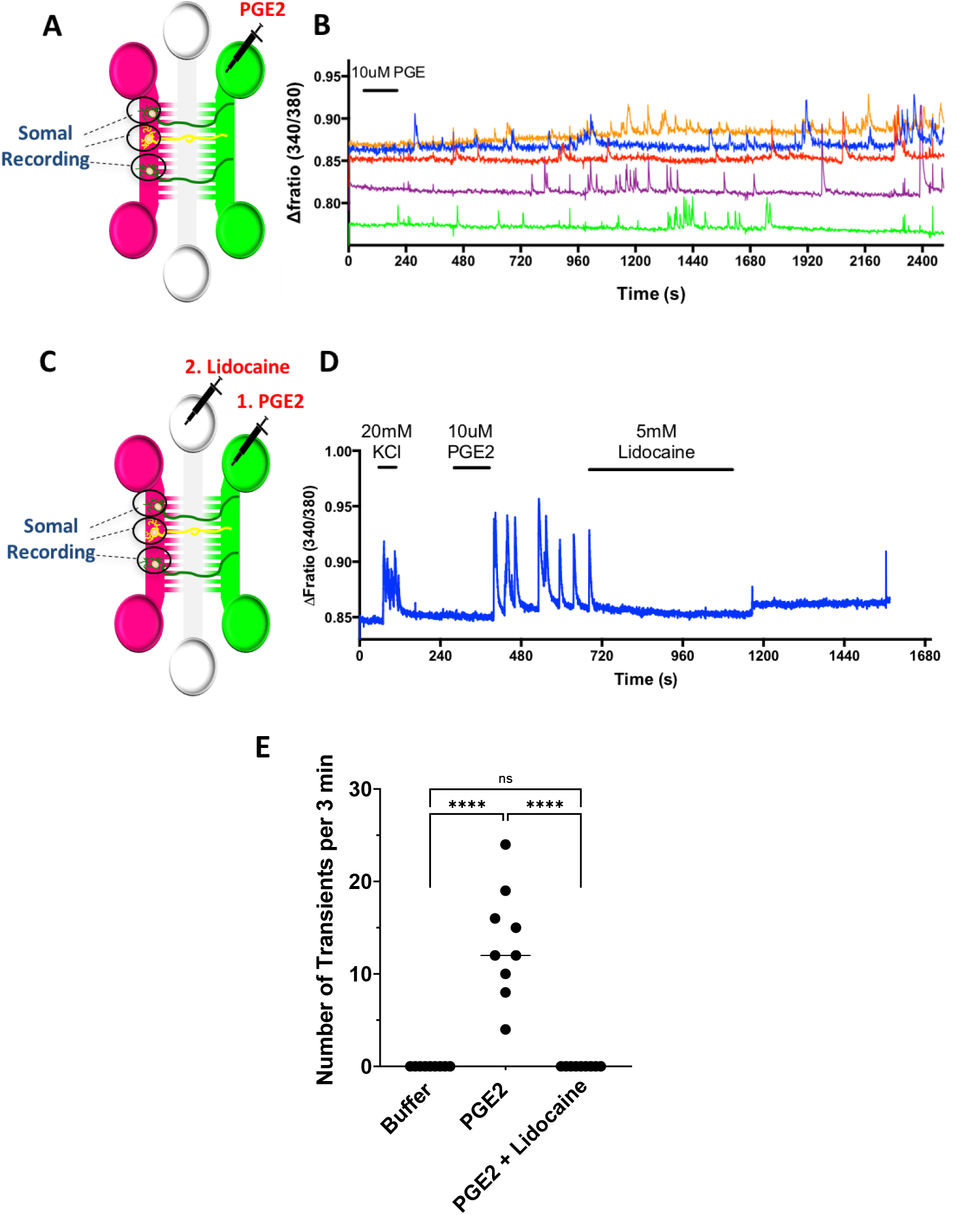
PGE2 evokes a persistent axonal activity which is abolished by lidocaine. **A)** Schematic of experimental set up used for data shown in panels B, three-compartment MFC is depicted with the somal compartment (pink), middle (white) and distal axonal compartments (green). **B)** Representative trace showing the Ca^2+^ transients recorded at the soma. Perfusion of 10μM PGE2 for two minutes resulted in axonal activity detected at the soma, which persisted after washout of PGE2. **C)** Schematic of the experimental set up used for data shown in panels **D** and **E**, using the three-compartment microfluidic configuration. KCl was applied to the distal axon compartment, followed by PGE2. 5mM lidocaine was then applied to the middle axonal compartment. **D)** Representative Fratio trace showing the calcium transients evoked by the application of PGE2 and blocked by the perfusion of 5mM lidocaine in the middle axonal compartment. **E)** Summary of the number of transients observed during a 3 minute window following stimulus application. An increase in the number of transients after application of PGE2 compared to baseline and a significant reduction in the number of transients after application of lidocaine was observed. (Kruskal–Wallis test with Dunn’s multiple comparisons test, ****p<0.0001, n = 9 from 4 cultures).

To localize the site of activation of the persistent activity, we examined the effect of the application of the anesthetic Lidocaine, a non-selective sodium and calcium channel blocker, to the central compartment **(Figure 2C)**. Pretreatment of the middle compartment with Lidocaine (5mM) completely and irreversibly blocked the PGE2 evoked persistent responses detected at the soma (**Figures 2D and E)**. This experiment demonstrates that the persistent responses elicited by PGE2 originate in the axon terminals.

We next examined whether cAMP-dependent signaling downstream of PGE2 receptor activation is required for the persistent activity observed in the axons. A three-minute application of Rp-cAMPS, a membrane permeant inhibitor of cAMP-dependent protein kinases, reversibly and completely blocked the PGE2-evoked persistent activity in the axons **(Figures 3A-C)**. Taken together, these data establish that the inflammatory mediator PGE2 evokes a local persistent activity in the DRG axons, and that this persistent activity requires the generation of cAMP and activation of PKA.

**Figure 3.**
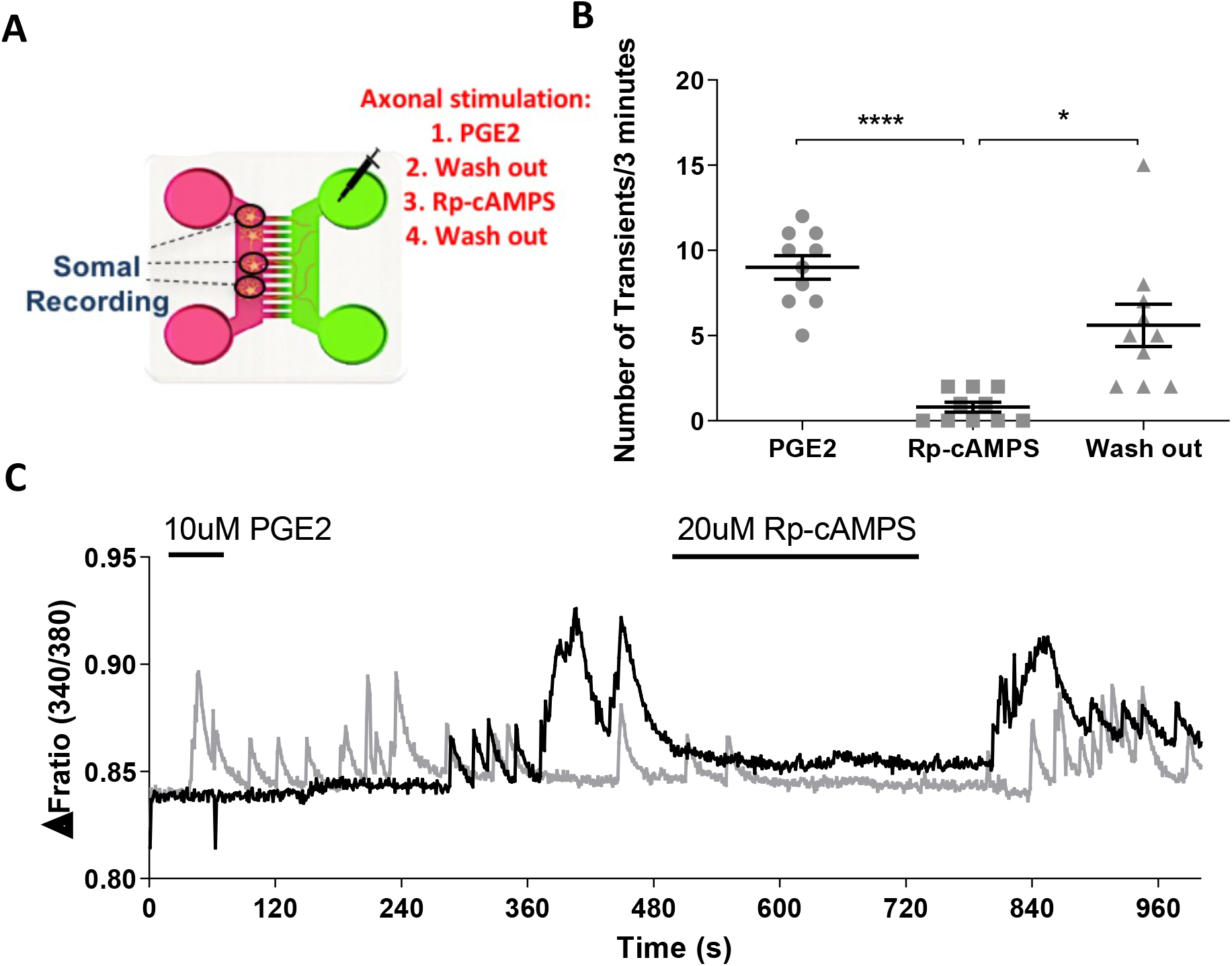
The PGE2-evoked persistent axonal firing is dependent on the cAMP/PKA signaling pathway. **A)** Schematic of the experimental protocol. Stimulation of the axonal side (pink) with PGE2 was followed by a wash and application of Rp-cAMPS. Ca imaging was performed from the somal compartment. **B)** Scatter plot quantifying the frequency of the Ca2+ transients observed. Rp-cAMP blocked PGE2-induced transients). The activity partially recovered following washout of Rp-cAMPS, **C)** Representative trace showing repetitive firing activity recorded at the soma in response to 1-minute axonal stimulation with PGE2. The activity is blocked with the application of 20µM Rp-cAMPS for 3 minutes. (One-way ANOVA with Tukey’s multiple comparisons test, *p<0.05, ****p<0.0001 n = 11 from 3 cultures)

### PGE2 application depolarizes axonal membrane in the absence of other stimuli

We hypothesized that the observed persistent activity, initiated locally at axonal endings and propagating to the soma is due to the depolarization of the axonal membrane in response to PGE2. Using the voltage-sensitive dye Fluovolt (*21, 22*), we measured the changes in axonal membrane potential in the axonal compartment (**Figures 4A-D)**. Application of KCl (30mM) for 90 seconds produced a change in fluorescent intensity of 5.18% ± 0.11% which returned to baseline immediately after wash with buffer **(Figure 4D)**. Application of PGE2 (10μM) produced a robust 5.89% ± 0.09% increase in the FluoVolt fluorescence intensity (**Figure 4E-G)**. The PGE2 evoked depolarization exhibited slow activation kinetics and a sustained plateau following the wash **(Figure 4E)**, supporting the activation of a second messenger pathway that could induces this persistent depolarization. Altogether these data demonstrate a direct and local depolarization of the axonal membrane in response to PGE2.

**Figure 4.**
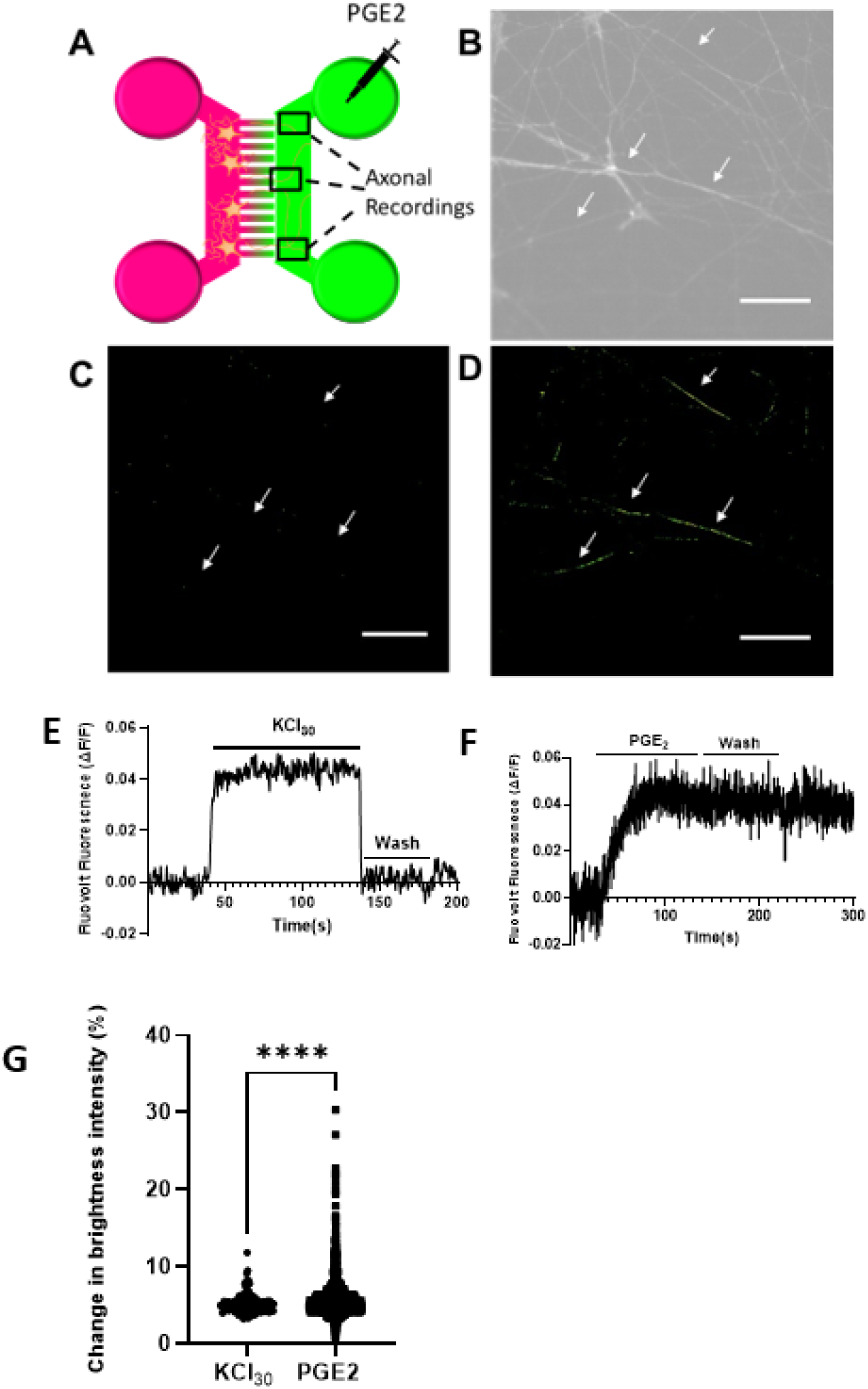
Application of PGE2 raises the membrane potential in the axons. **A)** Schematic of experimental set up for data shown in panels **B-G**. A two-compartment microfluidic configuration was used, where PGE2 or KCl were applied in the axonal side (pink). FluoVolt fluorescence intensity was measured in the axonal compartment. **B)** Bright field image showing a representative field of view (FoV) from the axonal compartment of a microfluidic culture. **C)** Background subtracted image (ΔF/F) of the FoV shown in **B**, prior to PGE2 or KCl application. No change in brightness intensity occurs prior to PGE2 application. **D)** Background subtracted image (ΔF/F) of the FoV shown in **B**, after PGE2 application. Changes in FluoVolt brightness intensity upon PGE2 stimulation are highlight with arrows. **E and F)** Representative traces of FluoVolt brightness intensity from the axonal compartment showing increased intensity with 30mM of KCl (E) or 10μM PGE2 (F). The KCl trace returns to baseline after washout whereas the PGE2 trace remains elevated. **G)** Summary of the data describing peak KCl and PGE2 evoked changes in FluoVolt fluorescence. Application of 10 uM PGE2 to the axonal compartment elicited a 5.89% ± 0.09% increase in fluorescent intensity comparable to the 5.18% ± 0.11% increase evoked by 30 mM KCl. (Student’s t-test, p<0.354, n_ROIs_= 925 on 20 devices from 18 cultures for PGE2 and n_ROIs_=135 on 4 devices from 3 cultures for KCl)

### PGE2-evoked persistent activity in sensory axons requires Nav1.8 channel activity

Given the observed silencing of PGE2-evoked axonal activity by the application of lidocaine to the middle chamber, which suggested a likely block of the propagation of the generated action potentials to the soma, we next considered the role of voltage-gated sodium channels in the generation of the persistent activity evoked by PGE2 in the axons. We found that application of 500nM TTX did not attenuate the PGE2 evoked persistent activity in the axons **(Figures 5A-C)**, suggesting that TTX-sensitive sodium channels are not involved in the generation or propagation of this persistent activity. We further examined whether the blockade of the TTX-sensitive Nav1.7 channels would affect the PGE2-mediated depolarization of the axons. We stimulated sensory axons with PGE2 for 90 seconds in the presence or absence of a selective Nav1.7 channel blocker P-05089771 (8nM) and monitored changes in axonal membrane potential using FluoVolt **(Figure 5D)**. We assessed the effect of the Nav1.7 blocker on membrane potential in two ways. We quantified the proportion of axonal area where the FluoVolt signal was increased by PGE2 stimulation (see Methods) either alone or in the presence of Nav1.7 blockers. We observed no significant change in the proportion of axonal membrane that exhibited a change in membrane potential, in response to PGE2 in the presence or absence of the Nav1.7 blocker **(Figure 5E)**. We observed however a slight decrease trend in the average change in the membrane potential in the axons in the presence of the Nav1.7 blocker **(Figure 5F)** suggesting a contribution from Nav1.7 channels to the generation of PGE2 evoked depolarization.

**Figure 5.**
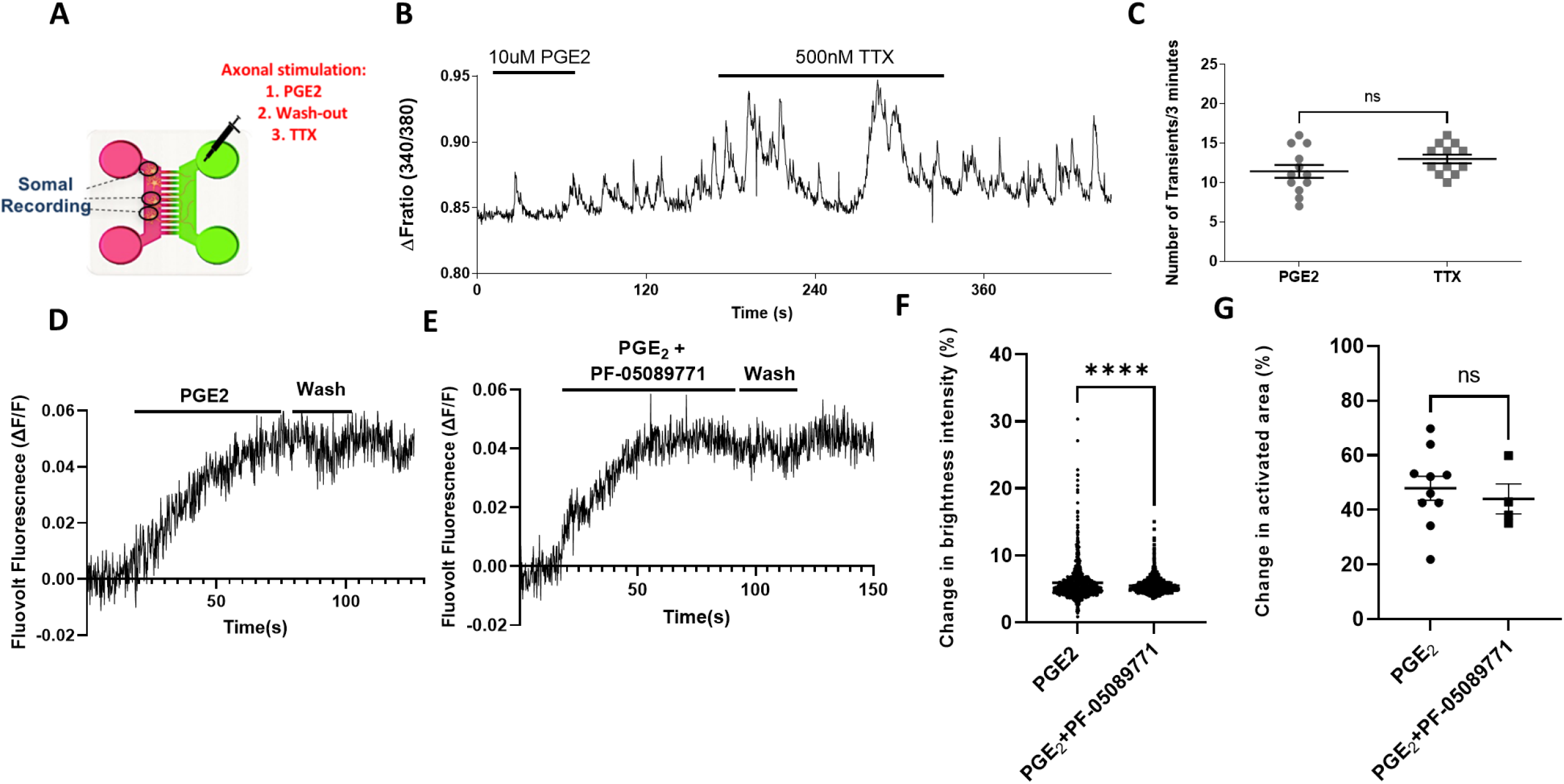
The PGE2-evoked persistent activity in the axons is not dependent on TTX-sensitive sodium channels. A) Schematic of the experimental protocol for data shown in panels B and C. Stimulation of the axonal side (pink) with PGE2 was followed by a wash-out and application of 500nM TTX while recording from the somal compartment. **B)** Representative trace showing repetitive firing activity recorded from the soma in response to a 1-minute application of PGE2 to the axonal compartment, which is maintained in the presence of 500nM TTX. **C)** Summary of the TTX effect on the number of transients observed after application of each stimulus. No significant change in the number of PGE2-induced transients after application of 500nM TTX was observed. (p=0.1261, One-way ANOVA with Tukey’s multiple comparisons test, n=15 from 3 cultures). **D and E)** Representative traces of FluoVolt brightness intensity in the axons showing the increase following stimulation with 10µM PGE2 (D) and 10uM of PGE2 followed by application of 8nM the Nav1.7 blocker PF-05089771. **F)** Summary of the data in D and E. Application of PGE2+Nav1.7 blocker produces a slight but significant decrease in the average peak brightness intensity (5.47 ± 0.03%, ****p<0.0001, Student’s t-test, n_ROIs_=925 on 20 devices from18 cultures for PGE2, and n_ROIs_=1202 on6 devices from 4 cultures for PGE2+ PF-05089771). **G)** Summary of the data showing the mean total PGE2 activated axonal area and PGE2 and the effect of PF-05089771. PGE2 application elicits activation of 47.94 ± 4.39% of the total axonal area, and addition of PF-05089771 with PGE2 does not significantly change the total activated area (44.03 ± 5.53%, p<0.5975 Student’s t-test, n=10 devices from 10 cultures for PGE2, and n=6 devices from 4 cultures for PGE2+ PF-05089771).

Nav1.8 channels have been shown to be modulated by PGE2 (*23*). We next considered the role of Nav1.8 channel in the PGE2-induced axonal depolarization. Using the three-compartment microfluidic configuration (Figure 5A), we found that application of the selective Nav1.8 blocker PF-01247324 (2nM) significantly attenuated the PGE2-evoked activity in the sensory axons (**Figures 6B and 6C)**.

**FIGURE 6.**
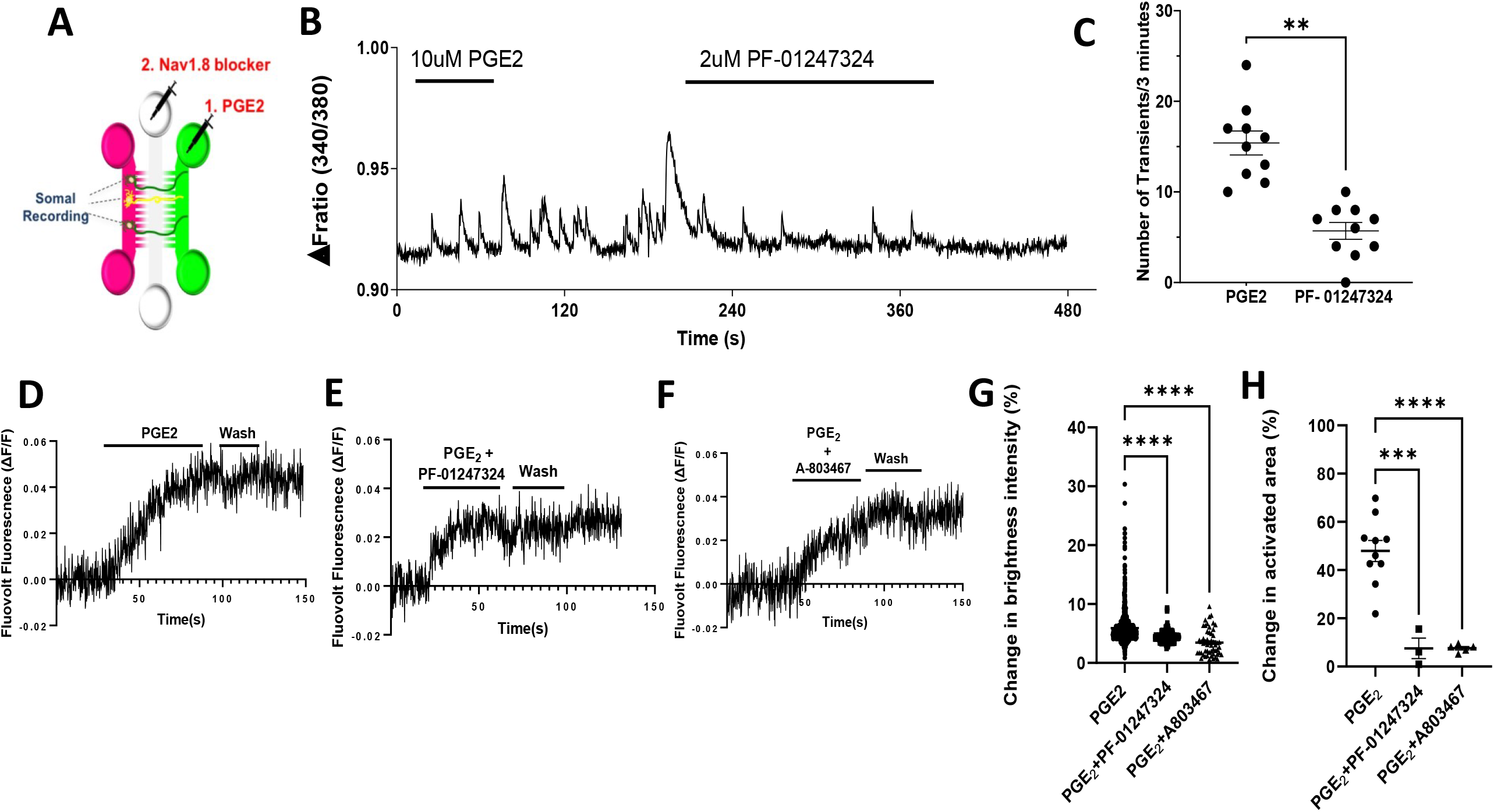
PGE2-evoked persistent axonal firing is predominantly mediated by Nav1.8 channels. A) Schematic of the experimental set up for data shown in B and C. Stimulation of the axonal side (pink) with PGE2 was followed by a washout and application of PF-01247324 while recording from the somal compartment. **B)** Representative trace showing repetitive persistent activity recorded from the soma in response to axonal stimulation with PGE2. Application of 2µM PF-01247324 largely attenuated the response. **C)** Summary of the data showing the effect of PF-05089771 on number of PGE2-evoked transients. A 3-fold decrease in the number of PGE2-induced transients after application of PF-01247324 was observed (**p<0.01, Wilcoxon matched-pairs signed rank test, n =10 devices from 3 cultures). **D-F)** Representative traces of FluoVolt intensity showing changes following treatment of the axonal compartment with PGE2, PGE2 with 2 nM PF-01247324, and PGE2 in the presence of 10 nM A-803467. Nav1.8 blockers attenuate the amplitude of the membrane depolarization evoked by PGE2. **G)** Summary of the changes in the brightness intensity evoked by PGE2. Application of PGE2 in the presence of the Nav1.8 blockers significantly reduces the average amplitudes of responses to nearly half (4.40 ± 0.035% for PF-01257324 and 3.46 ± 0.31% SEM for A-803467, **** p<0.0001, Brown-Forsythe and Welch ANOVA tests, nROIs= 518on 3 devices from 3 cultures for PGE2+PF-01247324, and nROIs= 53 on 5 devices from 5 cultures for PGE2+A-803467). **H)** Summary of the changes in total PGE2-evoked activated axonal area in the presence of Nav1.8 blockers. PGE2 application depolarizes 47.94 ± 4.39% of the total axonal area while Nav1.8 blockers significantly reduce the activated area. (7.53 ± 4.25% for PF-01257324 and 7.60 ± 0.73% for A-803467, ***p<0.0006 and **** p<0.0001 respectively, Brown-Forsythe and Welch ANOVA tests, n=3 devices from 3 cultures for PGE2+PF-01247324 and n=5 devices from 5 cultures for PGE2+A-803467 blocker).

We next performed voltage imaging experiments to determine the contribution of Nav1.8 channels to PGE2-evoked depolarization of axonal membrane. Using a two-compartment microfluidic configuration we examined the effects of two Nav1.8-selective channel blockers, PF-01247324 and A-803467 (**Figures 6D and E)**. While PGE2 evoked a 5.89 ± 0.09% change in FluoVolt fluorescence, co-application of PF-01247324 and A-803467 led to a statistically significant reduction in the observed change in the fluorescence to 4.40 ± 0.035% and 3.46 ± 0.31% respectively (**Figure 6F and G)**. When we measured the proportion of axonal area where the FluoVolt signal was increased by PGE2 stimulation, in the presence of the Nav1.8 blockers, we observed an almost complete abolishment (85% reduction) of PGE2-evoked axonal depolarization by the Nav1.8 blockers (**Figure 6H)**. For PF-01257324 only 7.53% ± 4.25 and for A-803467 7.60± 0.73% SEM VALUE of the total axonal area responded to PGE2-stimulation, while application of PGE2 alone led to 47.94% ± 4.39% of the axonal area exhibiting a change in fluorescence.

We also examined the potential contribution of HCN and ANO1 channels to the PGE2-evoked depolarization, and although application of zatebradine and ANO1 blockers zatebradine and T16-inh both led to reduced PGE2-evoked membrane depolarization in the axons, the results did not reach statistical significance (**Figure S5**). Altogether, these data strongly confirm the major contribution of Nav1.8 channels to the PGE2-evoked depolarization of axonal membrane.

### PGE2-induced depolarization is mediated by EP4 receptors

To further explore the mechanism of PGE2 evoked activation of sensory axons, we examined the role of the high-affinity prostaglandin receptor, EP4, in initiating the depolarization of the axonal membrane. We found that EP4 receptors are colocalized with Nav1.8 channels on DRG axons in microfluidic cultures (**Figure 7A, B and E)**. We observed a co-expression pattern of EP4 and the Nav1.8 channel along the adult DRG axons and EP4 staining was evident in a subset of Nav1.8 positive neurons (**Figure 7E)**. Next, we performed voltage imaging experiments on axons following a 20-minute incubation with the EP4 inhibitor L-161,982 (10nM). As shown in **Figure 7F**, the EP4 receptor inhibitor completely abolished PGE2-evoked axonal membrane depolarizations. The total axonal area activated by PGE2-stimulation in the presence of the EP4 inhibitor was 5.92 ± 1.92% of the total axonal area, as compared to 47.94% ± 4.39% of total axonal area that was activated by PGE2 alone. We also found a significant attenuation of the change in the amplitude of FluoVolt fluorescence, to 3.49 ± 0.085% by PGE2 in the responding axonal areas (**Figure 7G and H**). Therefore, this data supports the sole dependence of PGE2 on the high-affinity EP4 receptor for the generation of the persistent activity observed in the DRG axons.

**Figure 7.**
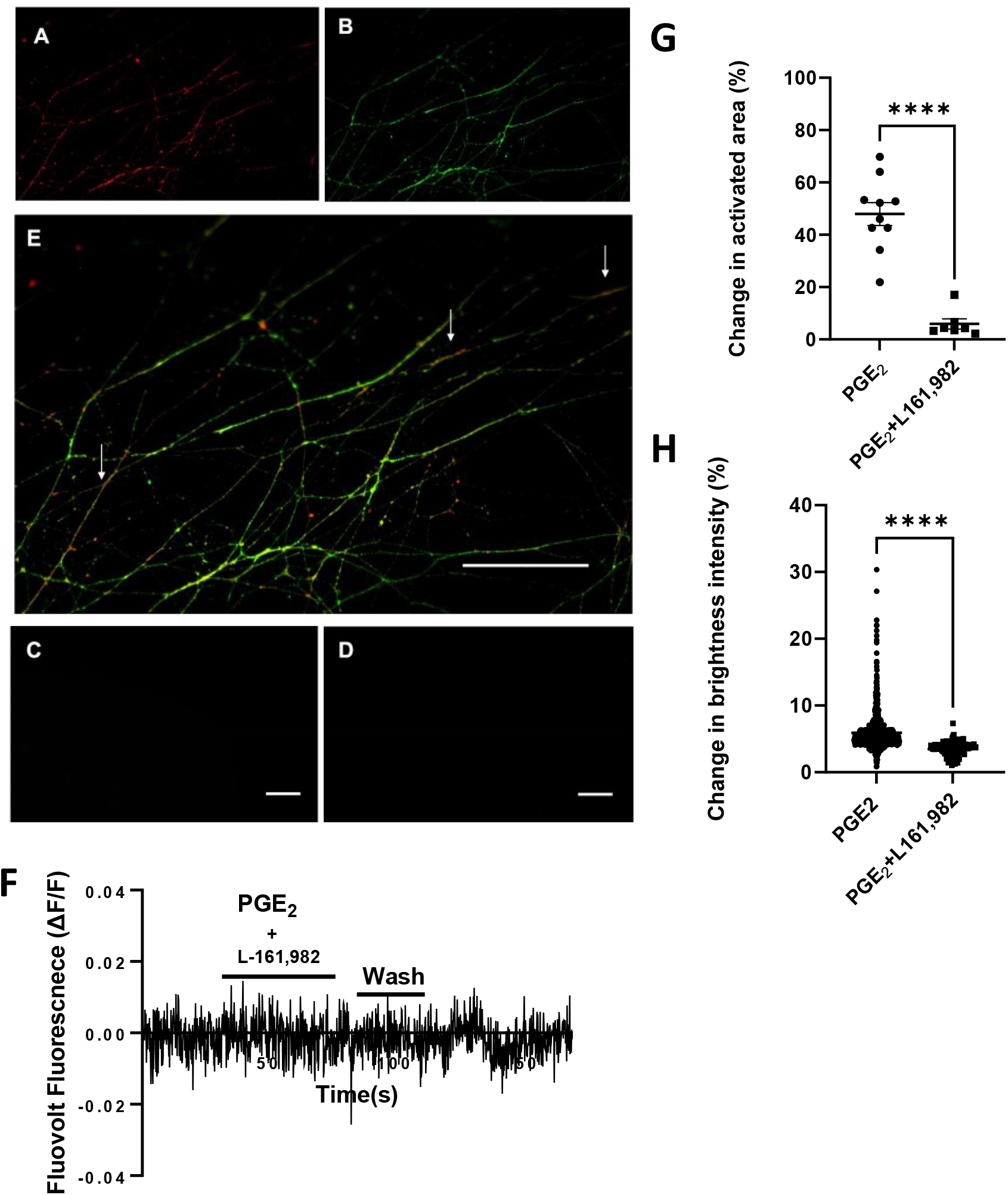
EP4 receptors colocalize with Nav1.8 channels in the axons and inhibition of EP4 receptors abolishes PGE2-evoked axonal depolarization. Representative images of adult DRG axons taken at day 5 in culture. **A and B)** Expression of EP4 in DRG axons in the MFC axonal compartment B) Na_v_1.8 staining in same FoV as A. **C and D**) Control for the secondary antibodies. **E)** An expanded view of an overlay image of EP4(red) and Na_v_1.8(green) staining from A and B, Arrows mark representative areas where co-localisation of EP4 receptors and Na_v_1.8 channels. Scale bars, 100μm. **F)** Representative FluoVolt intensity trace from an axonal area following a one-minute stimulation with 10µM PGE2 in the presence of 10 nM L-161,982. PGE2 failed to evoked a change in membrane potential in the presence of the EP4 inhibitor **G)** Summary of the changes in total PGE2 activated axonal area in the presence of L-161,982. L-161,982 significantly reduced mean total PGE2 activated axonal area (**** p<0.0001, Brown-Forsythe and Welch ANOVA tests, n=10 devices from10 for PGE2 and n=7 devices from 4 cultures for PGE2+L-161,982). **H)** Summary data describing the changes in the peak brightness intensity in the activated areas for PGE2-treated and PGE2+L-161,982 treated axons. Application of PGE2 in the presence of the selective EP4 inhibitor significantly attenuated the mean amplitudes (**** p<0.0001, Student’s t-test, nROIs=925on 20 devices from18 cultures for PGE2 and n_ROIs_= 141 on 7 devices from 4 cultures for PGE2+L-161,982).

## DISCUSSION

Taking advantage of a microfluidic culture model, we investigated sensitization of nociceptive axons downstream of PGE2/EP4 pathway. We have characterized the utility of the microfluidic culture platform for studying axonal physiology **(Tsantoulas et al, 2013)**. While previous *in vitro* studies have focused on events occurring at the cell bodies of DRG neurons, it is the nerve endings at the site of inflammation that are the first targets of inflammatory mediators. The fluidic isolation of axons in microfluidic cultures provides a more precise anatomical recapitulation of the peripheral neurons, and in addition the axons represent a closer physiological model of the nerve endings than the cell soma (*17*).

Using the microfluidic culture platform, we demonstrate that treatment of fluidically isolated DRG axons to PGE2 leads to a robust enhancement of the local responses to depolarization by KCl. Prostaglandins have been shown to target many channels in the membrane leading to enhancement of their responses. Previous studies for instance have shown the PGE2 mediated sensitization of ion channels such as TRPV1 in DRG cell bodies in culture (*24*). Nevertheless, stimulation by heat, mechanical or chemical stimuli ultimately leads to membrane depolarization and firing of action potentials in the nerve endings, hence sensitization of the axonal responses to depolarization described here could represent a common feature of inflammatory modulation independent of the processes that mediated enhancement of the local transducer channels such as TRPV1. We found that pharmacological blockade of HCN channels abolished the sensitization of KCl-evoked responses in the axons. HCN channels play an important role in regulating the firing frequency of DRG neurons and have been shown to be involved in neuropathic and inflammatory pain states (*19, 25, 26*). We found that blocking HCN channels with zatebradine led to a complete inhibition of PGE2-mediated sensitization of axonal responses to depolarization in our microfluidic model. This is consistent with the lack of PGE2-induced thermal hyperalgesia observed in mice with the HCN2 gene deleted from Nav1.8 expressing DRG neurons (*27*).

Intriguingly, we observed that PGE2 directly activated a subset of sensory axons leading to generation of persistent Ca signals recorded at the soma. We found that local activation of PGE2/EP4/cAMP/PKA pathway in the adult mouse DRG axons can consistently induce a membrane depolarization that results in the generation of action potentials propagating to the cell soma. Whilst the excitatory effects of bradykinin have been extensively characterized (*28, 29*), whether PGE2 can induce excitation of nociceptive nerve terminals is unclear. Previous studies have been inconclusive with some reporting that PGE2 failed to evoke spike discharge in primary visceral afferents and skin-saphenous nerve preparations (*30-33*), but that PGE2 can induce spike discharge in joint afferents (*34*). Species differences may account for the inconsistency in the *in vivo* observations, but technical limitations inherent to in vivo recordings (*17*) including conduction failure may also account for some of the negative results reported from *in vivo* studies. Studies on the excitatory action of PGE2 on sensory neurons in culture have primarily focused on the cell soma. For instance, application of PGE2 has been reported to evoke Ca^2+^ influx in dissociated rat DRG neurons (*35*), whereas, others have reported that exogenous application of the same concentration of PGE2 did not elicit any increase in the intracellular Ca^2+^ levels in DRG cell bodies (e.g. (*16*)). The utility of microfluidic cultures as a platform for the study of axonal properties overcomes many of the limitations of conventional cells cultures, as has been established by previous studies (*17, 36-39*). We have also demonstrated that the nociceptive axons in culture could provide a more physiologically relevant model of the nociceptive nerve endings(*17*). Several studies have examined the spatiotemporal dynamics of cAMP in the peripheral structures and have found a concentration gradient with the highest concentration of cAMP generated in the axons and the lowest in the soma during signaling (*40-43*). The differences in cAMP concentration can potentially explain the consistent responses of axons to the application of PGE2 in culture.

We hypothesized that the PGE2-evoked activity seen in the axons was the result of a sustained depolarization, downstream of cAMP/PKA activation. Using voltage imaging in the axonal compartment, we demonstrated that activation of EP4 receptor by PGE2, and the subsequent accumulation of cAMP lead to a local depolarization in isolated DRG axons. cAMP mediated depolarization of the membrane can occur through a number of mechanisms, including the closing of potassium channels, opening of Na+ or Ca-activated chloride channels. For instance, Vaughn and Gold have reported an increase in Ca-activated chloride currents and a decrease in potassium current in trigeminal afferents expose to an inflammatory cocktail including PGE2 (*44*). However, we found that pharmacological blockade of the calcium activated chloride channel ANO1 or of HCN channels however did not appreciably reduce the PGE2 evoked depolarization in mouse DRG axons. We subsequently focused on voltage-gated sodium channels, which possess a central role in pain sensation. PGE2 and cAMP have been shown to potentiate the TTX-R but not TTX-S currents in DRG neurons (*45-47*). We found that TTX-sensitive sodium channels are not mediating the depolarization induced by PGE2 in sensory axons, which is in keeping with the *in vivo* findings that deletion or block of Nav1.7 channels do not abolish peripheral sensitization induced by PGE2(*27, 48*).

Interestingly, blockers of the Nav1.8 TTX-R sodium channel significantly attenuated the depolarization induced by PGE2. It has been reported that acute administration of PGE2 can potentiate Nav1.8 current amplitudes as well as causing a hyperpolarizing shift in the current-voltage relationship (*47*). Further, the colocalization of Nav1.8 with EP4 receptors on nociceptive axons strongly implicatesNav1.8 as the main downstream mediator of PGE2-evoked depolarization in these axons. It remains unclear how changes in the magnitude of depolarization evoked currents or the small shifts observed in the current-voltage relationship could give rise to the significant depolarization observed in the axons in our experiments. It is possible that other channels are involved, resulting in subthreshold depolarization which is then amplified by Nav1.8 in the axons. For instance, potentiation of the heat sensor TRPV1 by PGE2 has been suggested to cause its activation by body temperature leading spontaneous or ongoing heat pain (*3*). A one-hour exposure to PGE2 has been reported to increases the surface expression levels of Nav1.8 via activation of PKA and phosphorylation of an RRR motif on the intracellular loop **(*49*)**. Interestingly, Kerr et al (2001) had reported that PGE2 mediated heat hyperalgesia was unaffected in Nav1.8 KO mice(*50*). However, compensatory mechanisms and sensitization of TRPV1 receptors could partly explain these results. The TTX-R sodium channel Nav1.9 has also been shown to play a role in inflammatory pain and is modulated by PGE2 **(*51*)**. The Nav1.8 blockers at the concentrations used in the present study, however, are not believed to interact with Nav1.9 channels. Altogether, the data presented here demonstrates a critical role played by Nav1.8 channels in mediating both inflammatory sensitization and also the direct excitation of nociceptive endings by PGE2, although the molecular mechanisms remain to be clarified.

In the present study we have validated the EP4/cAMP/PKA second messenger pathway leading to sensitization of responses to depolarising stimuli and generation of a sustained axonal excitation through activation of Nav1.8 channels in nociceptive axons. The persistent depolarization of nociceptive axons induced by PGE2 demonstrated here may potentially provide a general explanation for both acute and chronic manifestations of ongoing pain following tissue injury, independent of the sensitization of the specific receptor channels.

## MATERIALS AND METHODS

### Animals

Adult male C57BL/6 mice were purchased from Charles-River UK. All animals were maintained in a designated facility according to the UK Home Office Code of Practice for the Housing and Care of Animals Used in Scientific Procedures. Mice were sacrificed through CO_2_-anaesthetization followed by cervical dislocation in full compliance with UK Home Office Regulations and procedures under Schedule 1 of Animals (Scientific Procedures) Act 1986. All procedures were approved by the Animal Welfare and Ethical Review Body at King’s College London (PPL U136).

### Preparation of Microfluidic devices

The protocol used is a modification of the protocol described by **Tsantoulas et al**,**2013**. Microfluidic devices (Xona Microfluidics) with either 2 compartments (150 μm microgroove length) or 3 compartments (500 μm microgroove) were prepared according to published protocols (Vysokov 2016) Glass bottom dishes (WillCo Wells) were cleaned by sonication and sterilized with 70% ethanol. The dishes were previously coated with 0.5 mg/ml poly-L-lysine (Sigma). Microfluidic devices were non-plasma bonded onto the glass (Tsantoulas et al 2013). The assembled devices were coated after assembly with 40 µg/ml laminin (Sigma) dissolved in Neurobasal-A medium (Thermo Fisher).

### DRG cultures

Preparation of dissociated primary neuron cultures was performed through aseptic excision of the spinal column. Cervical, thoracic, lumbar and sacral DRG-containing cavities were exposed and DRG were collected. The ganglia were digested with enzyme mixture containing 0.125 mg/ml collagenase (Sigma) and 10 mg/ml dispase (Thermo Fisher Scientific) for 40 minutes in a 37 °C and 5% CO_2_ humidified incubator.

Following mechanical trituration, the dissociated neurons were centrifuged on a 10% bovine serum albumin cushion in HBSS. The cell pellet was washed twice and resuspended in complete medium containing Neurobasal-A (ThermoFisher), 1% B27 supplement (Invitrogen) and 1% Glutamax (Gibco), 100 units/ml penicillin and 100 μg/ml streptomycin.

The cells were loaded into prepared microfluidic devices and left to adhere for 60 minutes before being flooded with 200μl containing 150ng/ml NGF (Gibco) on the somal side and 130μl containing 200ng/ml NGF on the axonal side.

### DiI Tracing

The lipophilic trace dye DiI was used to mark cell bodies with axons crossing the microgrooves to the axonal side of the microfluidic devices. 24 – 48 hrs prior to the recording, the axonal compartments were incubated with the DiI solution (1:200 in complete medium medium) for 1 hour at 37 C. Care was taken to ensure microfluidic isolation remained intact. After washing with Neurobasal-A medium, and replacing with complete medium, the devices were placed in 37 C incubator until use. The stained cell bodies were visualized **(Figure S1)** immediately prior to live imaging on a Nikon TE200 microscope using a TRITC fluorescence filter set.

### Reagents

PGE2 stock (Sigma Aldrich) was dissolved in DMSO at 10 mM. Histamine dihydrochloride, Bradykinin and Serotonin (Tocris) stock solutions were prepared in DMSO. Tetrodotoxin (TTX) stock solution (Alomone Labs) was prepared in deionized H2O at 1 mM; Nav1.8 blockers PF-01247324, Nav1.7 blocker PF-05089771 (Tocris) were prepared in DMSO at 1mM; the non-selective sodium channel blocker lidocaine, the cAMP antagonist cAMPS-Rp, Nav1.8 blocker A-803467, EP4 inhibitor L-161,982, ANO1 blocker T16Ainh-A01, and the non-selective HCN channel blocker zatebradine hydrochloride (all purchased from Tocris) were dissolved in DMSO. All stock solutions were kept at -20 C until use.

### Drug perfusion

A bespoke perfusion system has been built for controlled pharmacological administration and avoidance of movement artefacts. The system is comprised of an 8-channel pen with a 360 μm removable tip (Automate Scientific) and regulated by a VC-8 Valve Controller (Warner Instruments) to allow 50 μl of solution every 5 seconds upon turning on.

### Calcium Imaging

After 4 days of culturing, both somal and axonal compartments were incubated with 2 μM of the calcium indicator dye Fura-2-AM (Invitrogen) in imaging buffer consisting of HBSS [Ca^2+^ and Mg^2+^-free] supplemented with 10 mM HEPES (ThermoFischer), 2 mM CaCl_2_ (Sigma), 1 mM MgCl_2_, 2 mM probenecid (Sigma), pH 7.4 with NaOH. Axonal and somal MFC compartments were washed twice with imaging buffer prior to recording. The devices were mounted on an inverted fluorescent microscope (Nikon Eclipse TE200) equipped with 20X Plan Fluor 0.5 NA objective and an EasyRatioPro imaging unit (Horiba Scientific). Images were acquired using ORCA 4.2 sCMOS camera (Hamamatsu). Bright field and DiI staining were visualized first and only neurons with staining in the soma (indicative of axonal crossing) were used for the analysis. The regions of interest (ROIs) were drawn around each corresponding soma and fluorescent images were acquired at 510nm at 6 Hz using 340 and 380 nm excitation wavelengths. Drugs were applied to the axonal compartment manually or using a perfusion system while recording from the somal compartment. Care was taken to keep the fluid level higher in the somal compartment to maintain fluidic isolation intact while performing the experiments.

### Voltage Imaging

For recording changes of in axonal membrane potential, a voltage sensitive probe, Fluovolt (Life Technologies) was employed. Cells were loaded with Fluo|Volt according to the manufactures recommended protocols (Life Technologies). Briefly, the axonal compartment was treated with 1:1000 dissolved Fluovolt in imaging buffer for 20 minutes at 37°C. After loading the axons were washed, and the devices were mounted on stage of the imaging microscope connected to EasyRatioPro. The excitation wavelength produced by the monochromator for fluorescent images is at 388 nm with an emission from the axonal compartment was recorded at 535nm at 6.3 Hz. Following stimulation by 30 mM KCl or 10 μM PGE2, axonal responses were recorded. Care was taken to preserve the fluidic isolation between somal and axonal compartments by maintaining the fluid level in the compartments.

### Image analysis

For calcium imaging experiments, the F_340_/F_380_ ratios (F_ratio_) were determined for designated regions of interest (ROIs) using EasyRatioPro software and the F_ratio_ traces were analysed using Clampfit 9.0 (Molecular Devices). The responses to stimulation were visually verified and confirmed if the maximum increase in F_ratio_ was at least three times greater than the standard deviations of the baseline calculated from at least 50s prior to drug application **(Figure S2)**. The peak response (ΔF_ratio_) was calculated as the maximum increase in fluorescence intensity above the baseline. In some experiments, 60 mM KCl was applied to the somal compartment to determine the number of neuronal responders. For PGE2 induced activity, each transient event was defined as increase in F_ratio_ with a peak that was at least three times greater the standard deviation of the baseline. Baselines were calculated from at least 10 seconds prior to application of stimulus. All events were visually inspected and confirmed as a response.

The detection of FluoVolt fluorescence intensity changes was defined as the (ΔF/F) between baseline responses taken at least 10 seconds prior to application of stimulus where no fluorescence activity was reported and ΔF maximum peaks within the timespan of application

For voltage imaging experiments, frame by frame TIFF files were transferred from EasyRatio Pro to FIJI, where axonal ROIs were selected according to brightness intensity changes. We quantified the changes in axonal membrane potential in microfluidic cultures by determining the average peak change in fluorescent intensity across all axonal regions in the field of view (FoV). The average peak change in fluorescence intensity was calculated by first determining a mean baseline image from 50 frames prior to drug application and subtracting the mean baseline image from all image stacks to create a ΔF image stack. The ΔF images were thresholded to demarcate the activated regions of in the axons. Particle Analysis in FIJI was used to assign the ROIs. The average peak in FluoVolt fluorescence intensity for each axonal ROI was calculated using the formula: 100 (Mean Peak Response-Mean Baseline Response) / Mean Baseline Response).

We also determined the proportion of the total axonal area that showed a change in FluoVolt fluorescence signal intensity in the FoV. As the limitations of FluoVolt did not allow longer recording times needed to determine the responses in the same FoV to the application of the inhibitors, we used the proportion of axonal area that showed an increase in FluoVolt signal to avoid any bias caused by the heterogeneity of the axonal responses between the experiments. The quantification of axonal areas that exhibited a change in membrane potential was represented as a percentage of the total axonal area that showed a change in FluoVolt fluorescence following drug application. Total axonal areas in each image, expressed as total number of pixels, were determined prior to drug application by thresholding the images in Fiji and selecting the entire axonal regions over the background. The activated axonal areas were determined from background subtracted images that were thresholded to highlight the areas that showed a change in intensity following drug application. The percentage ratio of the activated area to the total axonal area was taken as a measure of the activated area.

### Statistics

All data are reported as mean ± SEM unless otherwise stated. Statistical analysis has been performed using GraphPad Prism 9. Statistical significance was set at p < 0.05. Student t-test (unpaired, equal variance) was used to compare groups in calcium imaging analysis. One Way ANOVA with Tukey’s multiple comparison test was used for comparisons between three or more independent groups in calcium imaging analysis. Brown-Forsythe and Welch ANOVA tests were used to determine significance between groups of equal and different variance respectively on voltage imaging analysis.

## Supplementary Materials

Supplementary figures S1 to S5

## Acknowledgments

We would like to thank M. Malcangio and S.B. McMahon for their insightful comments on the project. We also thank N. Vysokov for helpful discussions and technical assistance.

## Author contributions

R.R., N.R., G.K., and P.A.M. designed the experiments. N.R., G.K., B.J.J. and R.R. carried out data collection and analysis. R.R. and P.A.M. supervised the study.

## Data and materials availability

All data discussed in the manuscript and required for the evaluation of the conclusions have been presented in the Results and in the supplementary figures. All data will be available upon request.

