## Supplemental Figures for "A microfluidic-based model of nociceptor sensitization reveals a direct activation of sensory axons by prostaglandin E2"

### Supplementary Figures

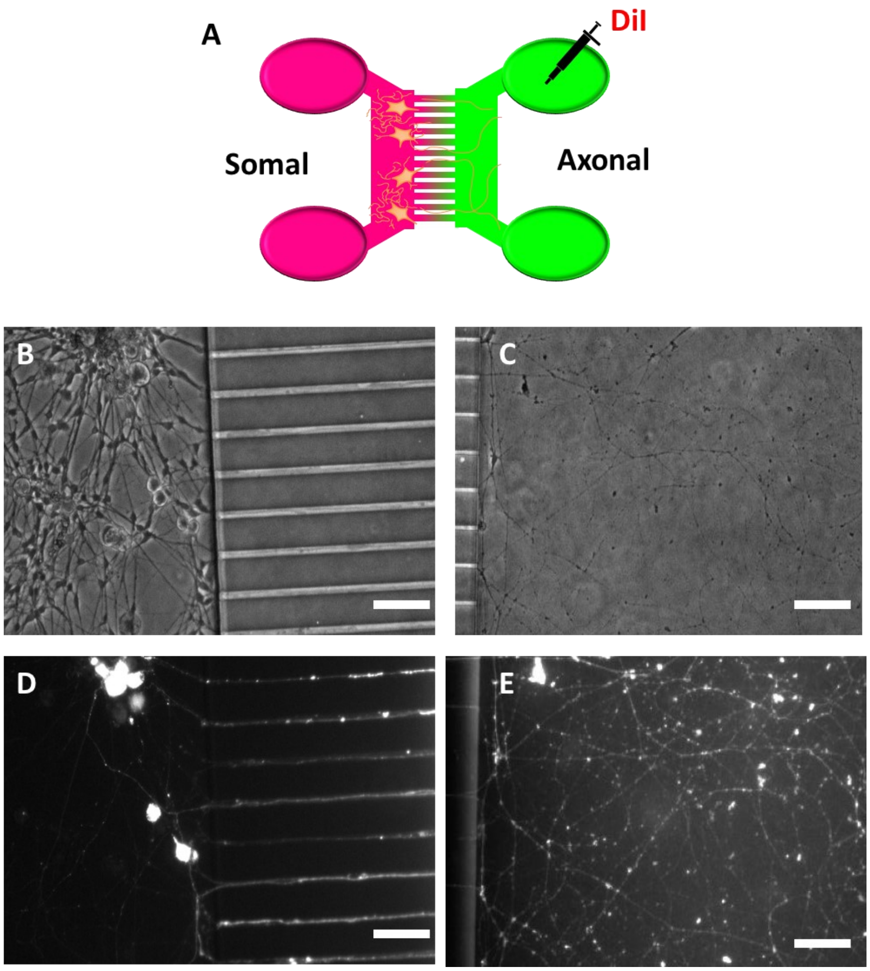


**Figure S1. Images of adult DRG neurons at Div 7 in MFC devices (20x objective). A)** Schematic of MFC culture setup, where the somal compartment is fluidically isolated from the axonal compartment where DiI is applied. **B)** Bright field (BF) image of the somal compartment, show the total number of DRG cell bodies in a given field. **C)** BF image of DRG axons shown in axonal compartment of MFC. **D)** DiI-stained DRG cell bodies in the somal compartment and their corresponding crossings towards the axonal compartment through the 450nM microgrooves. **E)** DiI-stained DRG axons in the axonal compartment. Scale bars, 100µm.

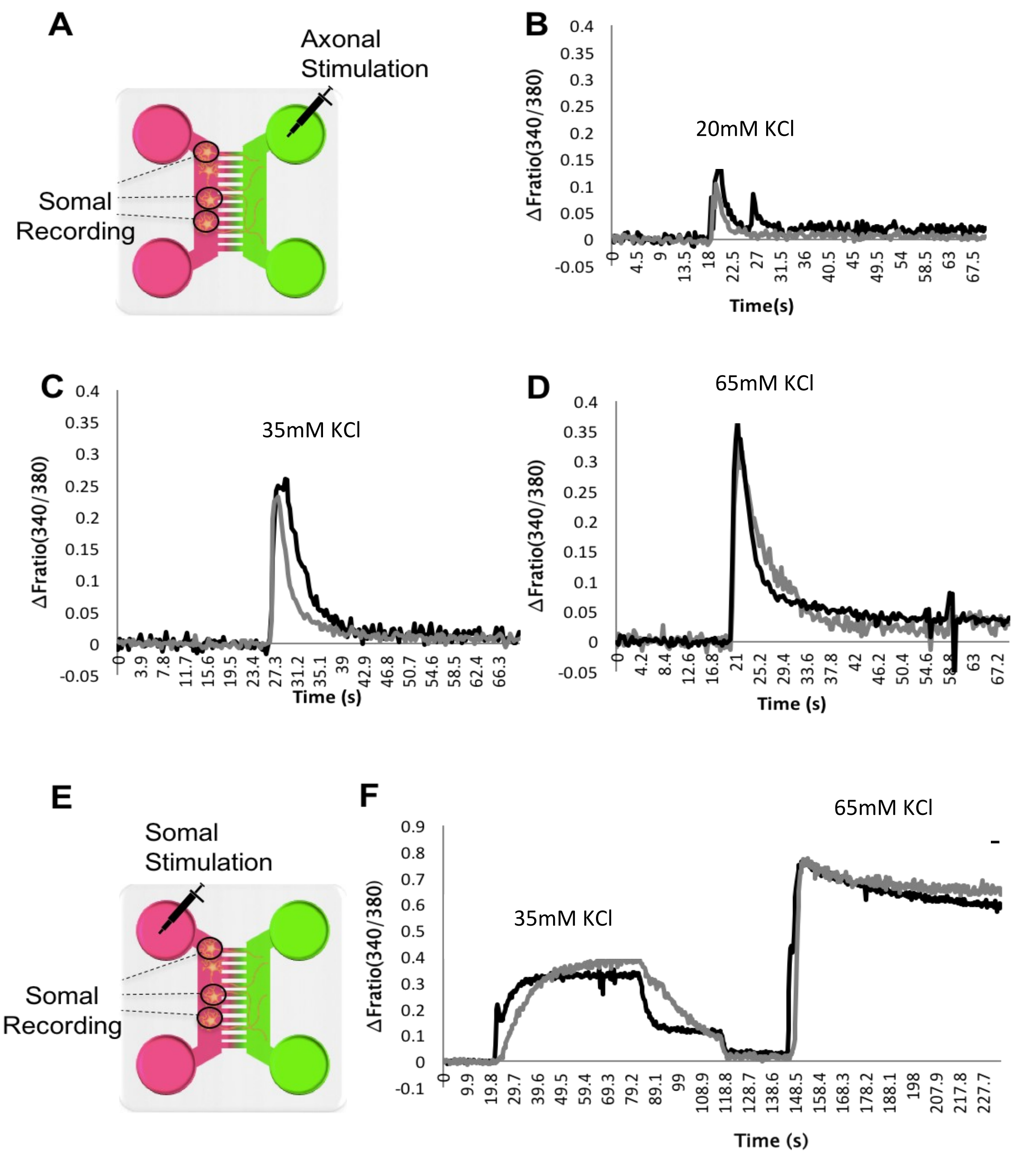


**Figure S2. Characterisation of axonal responses to chemical stimulation. Chemical stimulation of sensory neuron axons measured using Ca+2 imaging with readouts from the somal compartment.** Representative traces showing changes in calcium concentration at the soma in response to KCl stimulation in the axonal compartment. **A)** Schematic of experimental set up shown for data shown in panels **B-D)**. Upon chemical stimulation of the axonal compartment, changes in cell body $[Ca^{2+}]_{in}$ was measured. **B-D)** Representative traces showing changes in calcium concentration (measured as ΔF ratio), in response to axonal stimulation with 20mM,35mM and 65mM of KCl respectively. **E)** Schematic of experimental set up shown for data shown in panels **F)**. Where both stimulation and recordings of changes in $[Ca^{2+}]_{in}$ where measured in the somal compartment upon chemical stimulation. **G)** Representative traces showing changes in calcium concentration (measured as ΔF ratio) in response to somal stimulation with 35mM and 65mM KCl.

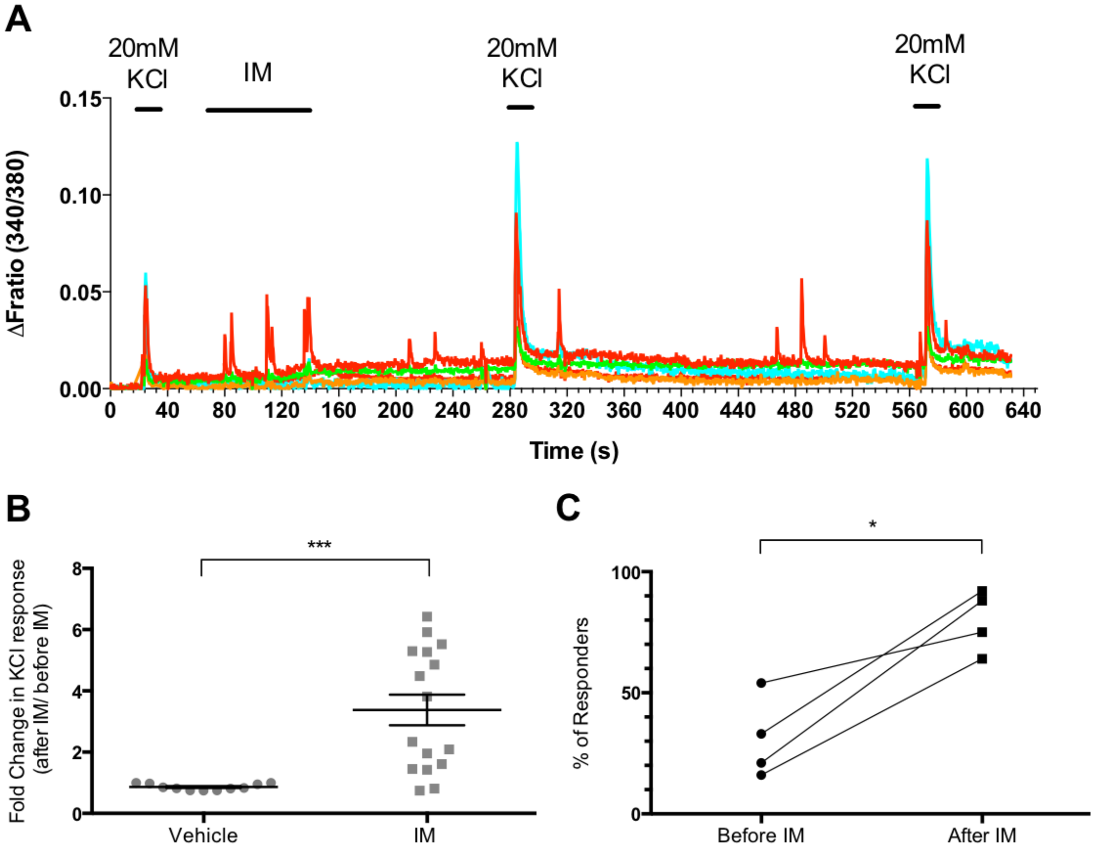


**Figure S3. Acute localised peripheral treatment of primary afferents with inflammatory mediators for two minutes resulted in two types of axonal sensitisation.** Acute localised axonal treatment with inflammatory mediators (IM), consisting of 10uM PGE2, 10uM Histamine, 10uM Bradykinin and 10uM Serotonin, resulted in two types of sensitisation of the axon. **A)** Representative trace showing axonal responses to first KCl stimulation (20 second), IM stimulation (2 minutes) and a potentiated response to the second KCl stimulation (20 seconds), which is maintained in the third response to KCl stimulation (20 seconds) after a 10-minute wash period. **B)** Fold change in response to KCl stimulation after vehicle treatment in comparison to IM treatment expressed as mean ± S.E.M. IM- treated neurons showed a much amplified peak response to KCl stimulation following IM administration, in comparison to vehicle treated neurons ***$p<0.001$, Student's t-test, n=20 cells from 4 cultures). **C)** Mean percentage of responder neurons in each culture. An increasing trend is observed in the number of DRG responders to KCl stimulation after two-minute IM treatment compared to KCl stimulation prior to IM treatment *$p<0.01$, Chi-square test, n=20 cells from 3 cultures).

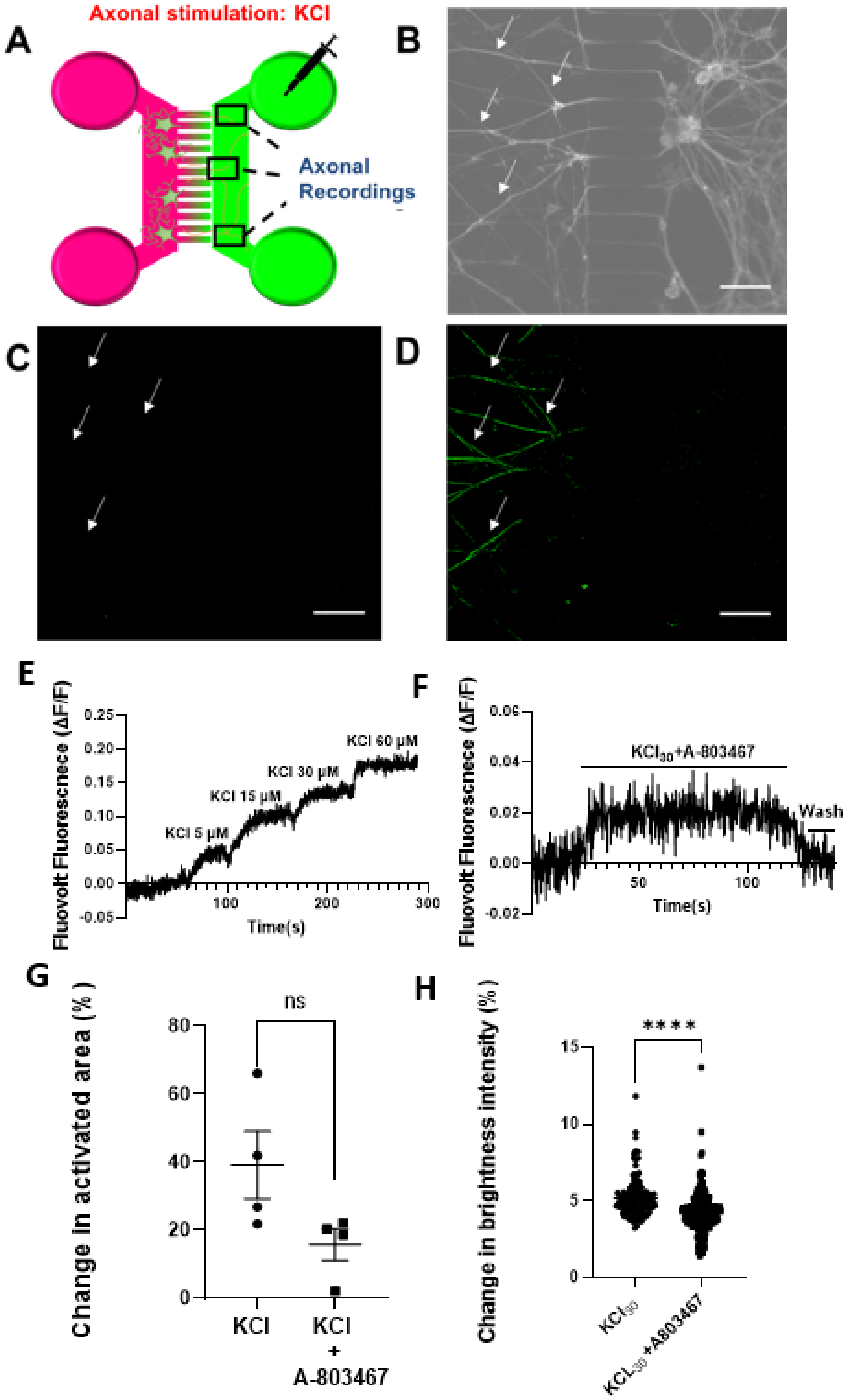
A
Axonal stimulation: KCl
Axonal
Recordings
B
C
D
E
Fluovolt Fluorescnece (ΔF/F)
KCl 5 μM
KCl 15 μM
KCl 30 μM
KCl 60 μM
Time(s)
F
Fluovolt Fluorescnece (ΔF/F)
KCl30+A-803467
Wash
Time(s)
G
Change in activated area (%)
ns
KCl
KCl
+
A-803467
H
Change in brightness intensity (%)
****
KCl30
KCL30+A803467

**Figure S4. KCl-induced depolarisation of the axolemma is detectable using Fluovolt dye A)** Schematic of experimental set up used for data shown in panels **B-D)**, 2-channel MFC compartment, where the somal side (pink) is separated from the axonal side (green). Following 1-minute application of 20mM KCl to the axonal compartment, changes in Fluovolt fluorescence were recorded from the axonal compartment. **B)** Bright field image showing chosen field of view (FoV) in the axonal compartment of the MFC. **C)** Background subtracted (ΔF/F) image of the same FoV as shown in B, before application of KCl. **D)** ΔF/F showing FoV after KCl application to the axonal compartment. White arrows highlighting examples of axons where changes in fluorescence intensity increased in response to stimulation with KCl. Scale bars, 100μm. **E)** Representative trace of a step depolarization experiment where increasing concentrations of 5mM, 15mM, 30mM and 60mM applied. Changes in brightness intensity are expressed as Fluovolt Fluorescence (ΔF/F), of one section of an axon in response to KCl application. **F)** Representative trace indicating changes in brightness intensity following a 1-minute stimulation with 10uM of PGE2 in the presence of 10 nM of a selectiveNav1.8 blocker (A-803467) at an axonal region of interest and expressed as Fluovolt Fluorescence (ΔF/F). The presence of the selective Nav1.8 blocker attenuates the amplitude of the membrane depolarization induced by KCl. **G)** Scatter plot describing changes in the number of total activated axonal area in the presence of 30mM KCl in the presence of the Nav1.8 blocker. The Nav1.8 blocker exhibits an attenuating trend towards KCl-mediated activity of the total area covered by sensory axons. ns $p<0.05$, Student's t-test, n=4,devices from 4 cultures for KCl and n=4 devices from 3 cultures for KCl+A-803467). **H)** Scatter plot describing changes in brightness intensity between KCl-treated and KCl+Nav1.8 blocker-treated axons. Application of 30mM KCl in the presence of the Nav1.8 blockers attenuates the average amplitudes of responses (5.16 ± 0.11% for KCl alone vs 4.16 ± 0.08% for KCl+ A-803467. (**** $p<0.0001$ Student's t-test, , $n_{ROIs}$=135 on4 devices from 4 cultures for KCl and ,$n_{ROIs}$= 271 on 4 devices from 3 cultures for KCl+A-803467).

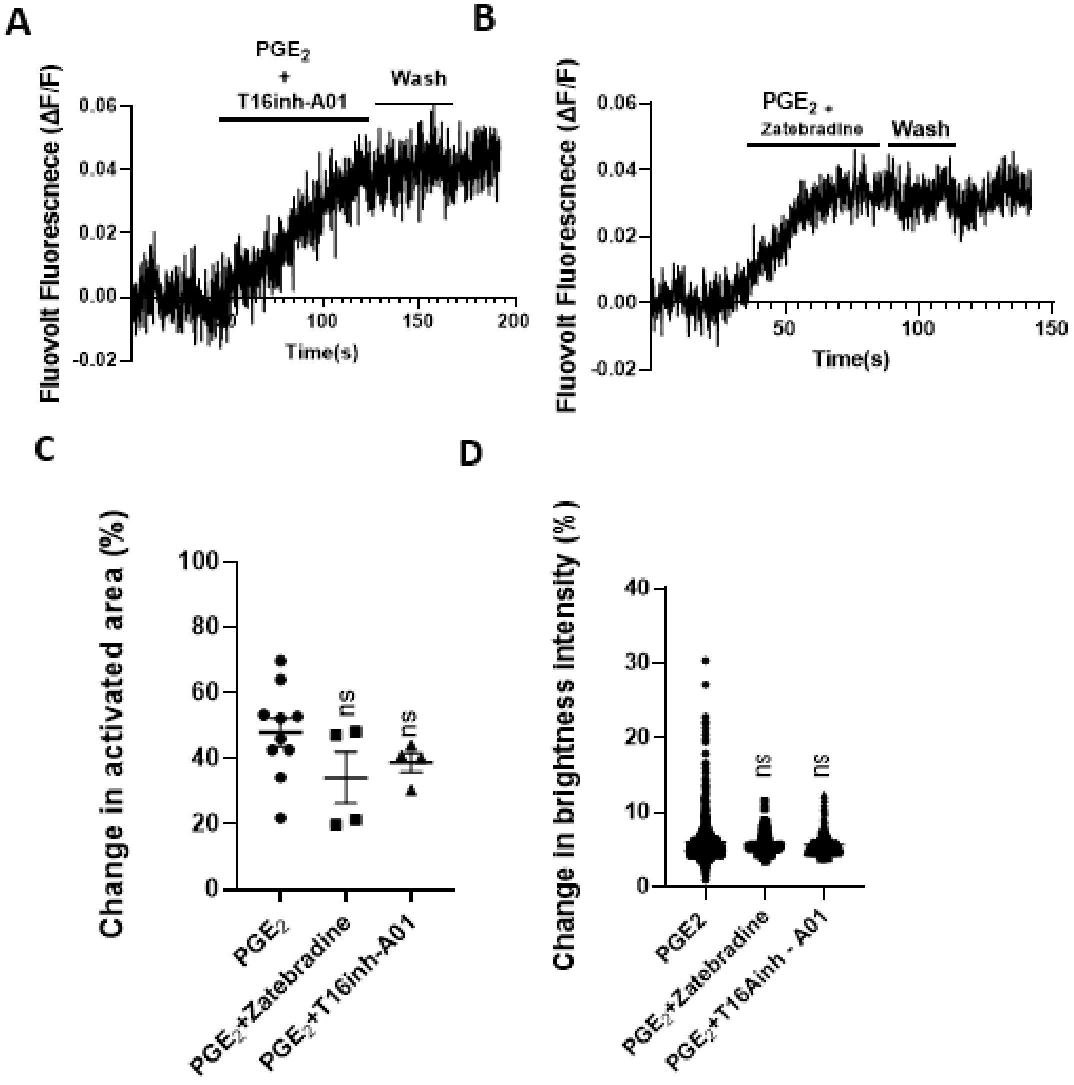


**Figure S5. PGE2-induced axonal depolarization is not mediated by ANO1 or HCN channels. A)** Representative trace indicating changes in fluorescence expressed as Fluovolt Fluorescence (ΔF/F), of an axonal region of interest in response to application of PGE2 in the presence of a selective ANO1 blocker (T16inh-A01). The kinetics and amplitude of the observed sustained depolarization remain unaltered despite the ANO1 ion channel blockade. **B)** Representative trace indicating changes in fluorescence expressed as Fluovolt Fluorescence (ΔF/F), of an axonal region of interest in response to application of PGE2 in the presence of a non-selective HCN channel blocker (Zatebradine). Zatebradine does not alter the amplitude or the kinetics of PGE2 elicited responses. **C)** Scatter plot describing changes in the number of total activated axonal area in the presence of 10μM PGE2 in the absence or presence of a selective ANO1 channel blocker and a non-selective HCN channel blocker in the presence of the Nav1.8 blocker. Neither blockers contribute significantly affected PGE2-activated sensory axons although a decreasing trend was observed. (ns $p>0.05$, Brown-Forsythe and Welch ANOVA tests, n= 4 devices from 4 cultures for PGE2+ANO1, n= 4 devices from 3 cultures for PGE2+HCN blockade). **D)** Scatter plot of average changes in Fluovolt Fluorescence between PGE2 and each ion channel blocker tested. PGE2 in the presence of the ANO1 and HCN blockers does not produce any significant effect in depolarization amplitudes generated by PGE2. p<ns for both (Brown-Forsythe and Welch ANOVA tests, , $n_{ROIs}$= 317 on 4 devices from 4 cultures for PGE2+ANO1 blockade and , $n_{ROIs}$=116 on 4 devices from 3 cultures for PGE2+HCN blockade).
